# Neuromechanical Phase Lags and Gait Adaptation in the Nematode *C. elegans*

**DOI:** 10.1101/2024.05.06.592744

**Authors:** Christopher J. Pierce, Yang Ding, Baxi Chong, Hang Lu, Daniel I. Goldman

**Affiliations:** School of Physics, Georgia Institute of Technology; School of Chemical and Biomolecular Engineering, Georgia Institute of Technology; Beijing Computational Science Research Center

## Abstract

Undulation is a locomotor strategy in which waves of bending propagate along the body.This form of locomotion is observed in organisms that span orders of magnitude in size and represent diverse habitats and species. Despite this diversity, common neuromechanical phenomena have been observed across biologically disparate undulators, due to common mechanics. For example, Neuromechanical Phase Lags (NPL), a phenomenon where waves of muscle contraction travel at different speeds than body bends, have been observed in fish, lamprey and lizards. Existing theoretical descriptions of this phenomenon implicate the role of physical body-environment interactions. However, systematic experimental variation of body-environment interactions and measurement of the corresponding phase lags has not been performed. Using the nematode *Caenorhabditis elegans* we measured muscle and body curvature dynamics simultaneously, performing calcium imaging in the body wall muscles while systematically varying the environmental rheology. A mechanical model demonstrates that the measured phase lags are controlled by the relative strength of elastic torques within the body and resistive forces within the medium. We further show that the phase lags correspond with a difference in the wavenumber of the muscle activity and curvature patterns. Hence, the environmental forces that create NPL also act as a filter that shapes and modulates the gait commanded by the nervous system. Beyond nematodes, the simplicity of our model further suggests that tuning body elasticity may serve as a general means of modulating the degree of mechanical control in other undulators.

## I. INTRODUCTION

Understanding how organisms coordinate their bodies for effective locomotion is challenging. Locomotion emerges from the dynamics of interacting hierarchical systems that span length scales from cells to tissues and neural networks to whole bodies and beyond: locomotion also necessarily involves non-trivial interactions between the body and the immediate physical surroundings. Body-environment interactions are an especially significant factor in organisms that rely on continuous body-environment contact for propulsion, such as snakes, fish, spermatozoa, and other lateral undulators. Lateral undulation involves the propagation of waves of body bending along typically elongated, slender bodies. It is observed in organisms found in both wet and dry environments, across orders of magnitude in size, and in both dissipative (low Reynolds number (Re) fluid[1] or granular material[2]) and inertial (high-Re fluid) regimes[3].

Despite the biological diversity among undulating organisms, and the inherent complexity and high dimensionality of their locomotory phenomena, emergently simple neuromechanical dynamics have been identified that are shared across diverse organisms and environment types. Common neuromechanics of undulation arise, in part, because of the centrality of body-environment interactions – common mechanical constraints often produce common control strategies. Correspondingly, mechanics often facilitates simple low-dimensional descriptions of neuromechanical phenomena that span biological diversity.

One common feature of undulation neuromechanics is the presence of so-called ‘neuromechanical phase lags’ (NPL)[4–6]. To propagate a bend along the body, a wave of muscle activation m(s, t) (where s is the body coordinate and t is time) is initiated and passed from head to tail. This wave of muscle activation and the resultant wave of body curvature κ(s, t), in general, do not travel at the same speed, producing a phase difference between m and κ that varies along the body. NPL were first experimentally observed in inertial aquatic swimmers[4], such as bass[7] and eels[4]. To explain the origin of the effect, detailed, high-dimensional models were developed, which incorporated large numbers of system-specific parameters, (such as body taper and muscle non-linearity)[8].

Subsequently, the effect was observed in low-inertia situations, such as the swimming of sandfish lizards in sand[2, 6], a frictional fluid. In this highly damped context, features of the low-inertia dynamics of the body-environment interactions enabled a low-dimensional description of the phenomenon[6]. The analytical tractability of the drag forces in this regime revealed that NPL could be understood simply as a consequence of torque balance, and therefore was likely a generic feature of undulation controlled by body-environment mechanics. However, these models were used to describe single organisms moving in a single environment – no systematic experimental variation of the body-environment interactions, and subsequent determination of the effect on the phase lags has been conducted.

This is likely the result of the experimental limitations of electromyogram (EMG) recordings typically used to measure NPL in free-moving organisms[2, 9]. Furthermore, the limited range of environments in which typical vertebrate undulators locomote, makes systematic variation of body-environment interactions challenging. Moreover, EMG recordings provide limited spatial resolution, due to the discreteness and invasiveness of EMG probes, limiting the ability to gather information about the spatial variation of NPL along the body. Overcoming these constraints will help to quantitatively connect body material properties and environmental rheologies to the observed phase lags, thereby explaining the ultimate neuromechanical origins of the phenomena.

We overcome these challenges by employing the nematode *Caenorhabditis elegans*, an invertebrate undulator and widely used model organism (see Fig. 1), which provides optical access to muscle activity through calcium imaging instead of invasive EMG probes. In the wild, *C. elegans* encounter complex heterogeneous terrain, such as the inside of rotting vegetative matter (Fig. 1) [10]. In the lab, they can locomote in a wide range of synthetic environments (fluids with viscosities spanning orders of magnitude[11, 12], viscoelastic fluids[13] granular media [14] and other heterogeneous terrains [15–17]. As environmental parameters (e.g. viscosity) change, they are known to adapt their gait parameters (e.g. wavenumber and frequency) in response to maintain performance[12]. This locomotory robustness permits systematic variation of the body-environment interactions that produce phase lags and direct testing of model predictions. Furthermore, it allows us to ask what role the phase lags play in selecting and coordinating the appropriate gait for a given environment. Because phase lags imply a difference in muscle and curvature wave speeds, they imply a difference between the wavelength commanded by the locomotor circuit and articulated by the muscles and the gait that results.

**FIG. 1.**
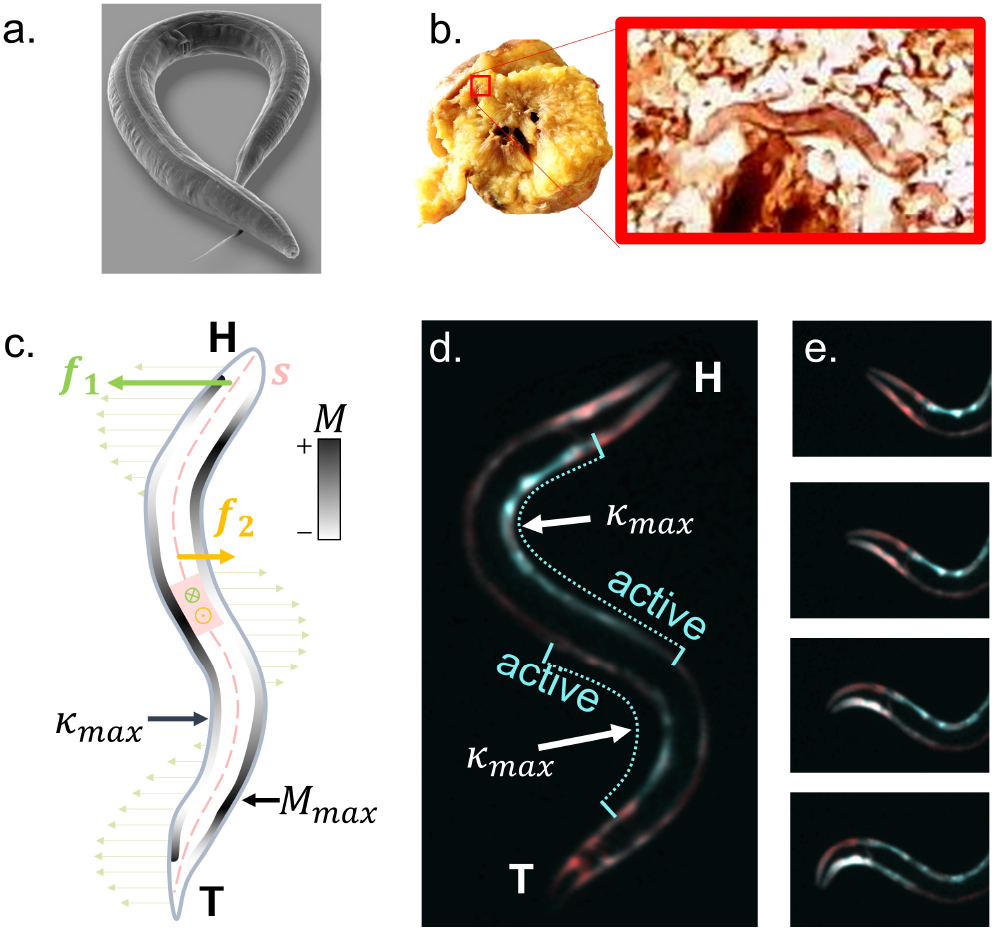
The nematode, *C. elegans* is a versatile undulator and provides optical access to neuronmuscular dynamics. a) a scanning electron microscopy image of *C. elegans* (courtesy of Ralf Sommer). b) a worm crawling through a thin slice of a rotten peach. c) schematic illustration of neuromechanical phase lags showing approximate environmental resistive forces, and torques acting on a point (red circle) from two points along the body (*f*_1_ and *f*_2_), regions of muscle activity and inactivity and local curvature. d) false color image showing the calcium-sensitive GCaMP3 fluorescence signal along with the calcium-insensitive RFP reference, illustrating approximate area of muscle activity/inactivity. e) time series showing increasing calcium signal on the left body wall muscles as a leftward head bend is initiated.

In this work, we measured body wall muscle activity and body curvature simultaneously for worms undulating in fluids spanning several orders of magnitude in viscosity, and on agarose pads. This allowed calculation of phase lags for a broad range of body-environment interaction regimes. We compare our results to a mathematical model based on the balance of external and internal torques and find good agreement, confirming that the magnitude and spatial variation of the phase lags are ultimately controlled by the relative importance of torques arising from within the body, and torques from the external environment, as suggested in [6]. Finally, we discuss the implications of the mechanical tuning of phase lags for locomotion control across environments. We show that the phase lags produce a corresponding change in the wavenumber of the muscle activity pattern relative to the curvature that is adaptive. As the worm actively adjusts its wavenumber to help maintain speed in more viscous environments, the same passive mechanical effects that produce the phase lags also modulate the commanded wave articulated by the muscles, mechanically enhancing the viscosity-induced wavenumber shift. In this way, we show that nematode gait adaptation is not a purely active process, but results from a combination of active sensory modulation of the gait and passive mechanical control.

## II. RESULTS

### A. Neuromechanical Phase Lags in *C. elegans* increase with viscosity

As described above, points of maximum curvature and points of maximum muscle activation do not necessarily coincide, leading to phase lags (See Fig. 1,c). This effect is predicted to generally depend on the relative strength of torques from external resistive forces, such as viscous drag, and internal body forces, which are primarily determined by the passive material properties of the body [6]. Previous nematode-specific neuromechanical models also predict phase lags to arise for worms crawling on agar pad, where drag forces dominate internal elasticity[13, 18, 19]. To test these models, we measured muscle activity patterns via ratiometric calcium imaging in the body wall muscles [20, 21] in *C. elegans* in a range of viscous fluids (buffer and 1−3% methylcellulose) and on agar surfaces. This allowed us to experimentally determine the effect of different internal/external torque ratios on the phase lags.

To perform calcium imaging, calcium-sensitive fluorophores, such as the family GCaMP proteins[22], are expressed in particular target cells or tissues. Their fluorescence intensity is proportional to the local calcium concentration, and therefore when expressed in muscles, indicates the degree of muscle activation [20]. However, because muscles necessarily contract and relax during movement, large changes in recorded intensity can also arise as the volume, and therefore the concentration of fluorophores, changes. Furthermore, motion artifacts and segmentation errors can lead to additional erroneous signals [23]. To eliminate these effects, typically a ratiometric approach [20, 22, 24] is used. A calcium-insensitive red fluorophore (RFP) is expressed in the body wall muscles along with a green calcium-sensitive GCaMP3, shown in Fig. 1, b,c and supplemental videos (Also see methods and [20]). By referencing the red, we can eliminate intensity variations that arise due to cell volume changes and movement. The measured ratio can therefore serve as a proxy for muscle activity, up to a constant time delay resulting from the protein kinetics [20, 22, 24].

To determine the phase lags, we first calculated the GCaMP-RFP signal ratio within the dorsal and ventral body wall muscles (See. Fig. 1) at each point along the body, along with the corresponding body curvature at each point, for all the environments. Fig. 2, a, b shows the ratiometric calcium signal for the left and right body wall muscles respectively and the curvature oscillation at the midpoint along the body for an example collected from buffer (Fig. 2,a) and crawling on agar (Fig. 2,b). The activity of the two body-wall muscles are approximately 180°out of phase as expected based on their contralaterally inhibited connectivity[25], but have an overall phase shift relative to the curvature oscillations, which varies along the body.

**FIG. 2.**
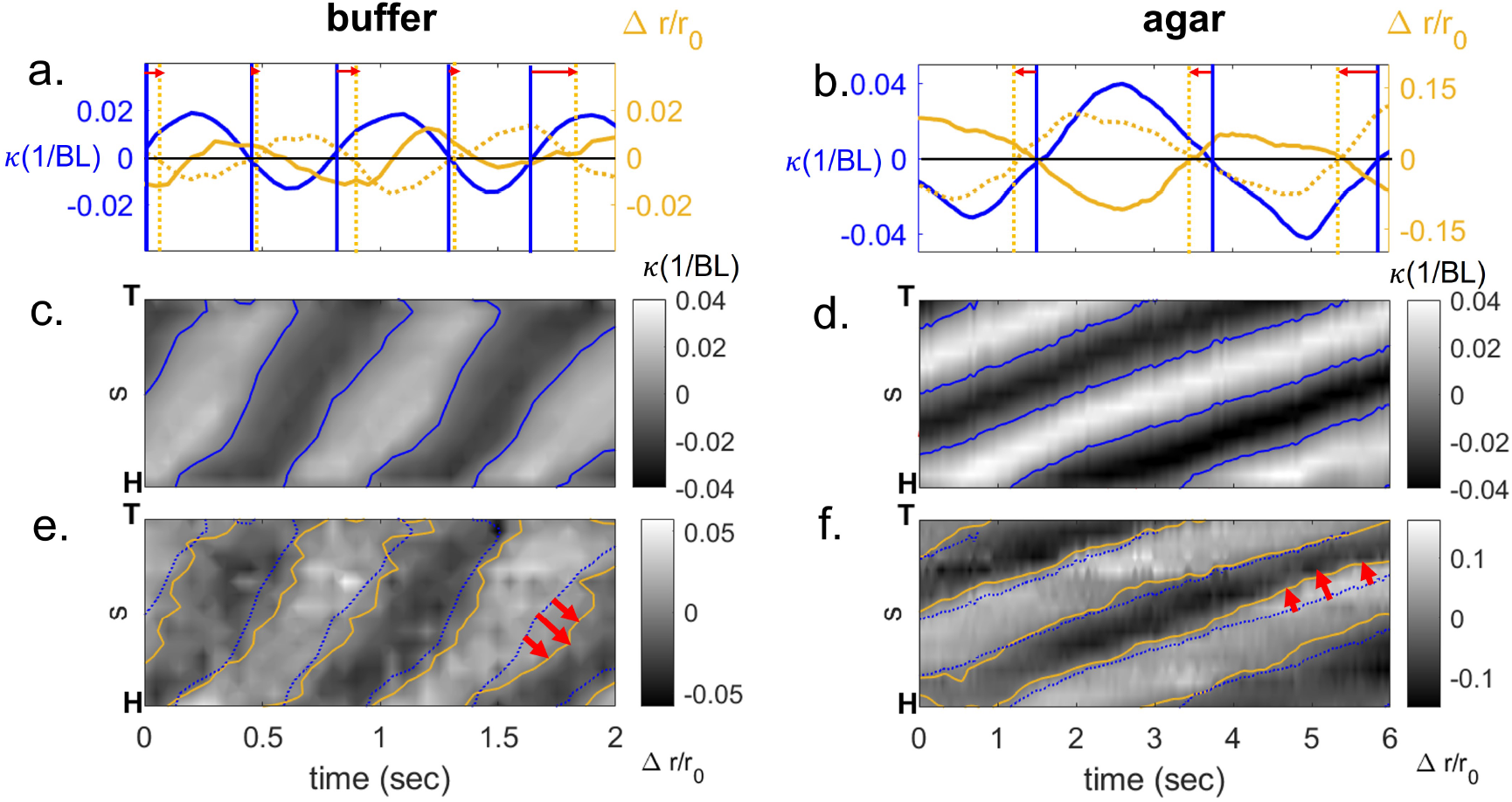
Comparing body curvature and muscle activation dynamics in buffer and agar. a) Ratiometric calcium signal from left (solid yellow line) and right (dotted yellow line) body wall muscles compared to body curvature oscillations at the midpoint for a worm swimming in buffer (a) and agar (b). Red arrows indicate delay between zero-crossings in curvature and muscle activity oscillations. Body curvature heatmap with nodelines highlighted in blue (c, d) and muscle activity heatmap (e,f) showing muscle nodelines (yellow) with curvature nodelines overlaid (in blue).

**FIG. 3.**
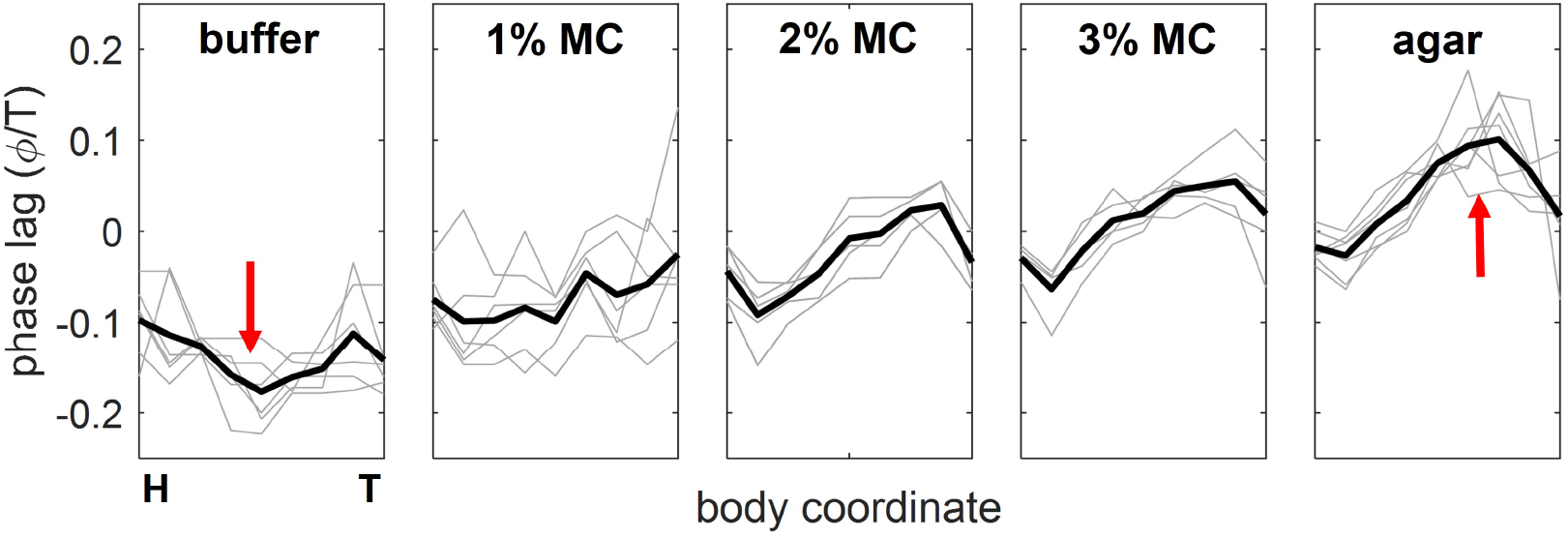
Neuromechanical phase lags for individual animals (gray lines) and population average (black lines) in a range of environments (buffer (N=6), 1% MC (N =6), 2%MC (N=5), 3% MC (N=5) and Agar (N=7)).

By computing analogous curves for a set of points extending from head to tail, we can construct space-time plots of muscle activation that can be compared with the corresponding curvature patterns, shown in Fig. 2,c-f. These results reveal differences in the overall shape of muscle and curvature waves in both buffer and agar, evident in the variations in the wave speed of each respective wave along the body, which can be visualized by comparing the slope of the line of zero crossings in muscle and curvature waves (blue and yellow curves in Fig. 2,e,f).

Neuromechanical phase lags were calculated from the spatiotemporal muscle/curvature data using a cross-correlation technique, which helped mitigate the effect of noise in the muscle signal (see methods). As noted above, the protein kinetics produce an overall delay of the GCaMP signal relative to muscle activation, which we have not taken into account. Hence, the absolute magnitude of the phase lags is ambiguous. However, as previously shown in [20], the delay from protein kinetics is consistent for a given environment, hence the *relative* phase lags of different parts of the body may be determined, with an additional delay which is constant along the body. In buffer, we observed a decrease in the relative phase lags from the head to the midsection (3), which returned toward zero in the posterior half of the body. Methylcellulose at 1% displayed nearly constant phase lags with a small increase along the body. As the wt-% was increased, and also on agar surfaces (thought to have higher overall resistive forces than the most viscous fluids considered here[13]), phase lags began to accumulate more significantly along the body. This phase lag accumulation displays an asymmetric spatial profile, increasing along the initial 70% of the body and then relaxing back toward phase alignment near the end of the organism (except for a small decrease in the phase lag near the head of the organism, which appears consistently across environments).

Taken together, our results suggest that in lower viscosity environments the relative phase lag across the body is approximately constant, or in the case of buffer, slightly decreasing in the mid-region. As the resistance in the environment is increased (by either increasing viscosity, or by switching to a solid gel surface), asymmetric phase lag accumulation along the posterior region increases in magnitude, in accord with prior qualitative predictions based on torque balance [6]. Our data also displays good agreement with a previous neuromechanical model of *C. elegans* from Denham et al [18, 19] and an earlier biomechanical model from Shen, et al [13].These models predict an approximately flat neuromechanical phase lag profile across the body for swimming in buffer, but for agar crawling predicts phase accumulation along the body.

Notably, most vertebrate systems appear to operate in phase-lag regimes similar to those observed for worms in high-resistance environments, displaying anterior-posterior phase lag accumulation. This suggests that they all operate in regimes where the external drag forces are the most significant determinant of the phase lags (relative to internal forces). Our results suggest that across environments, *C. elegans* interpolates smoothly between the different regimes predicted in [6]. Agar and high viscosity environments reproduce the spatial phase lag profiles of both dry, inertial sand-swimming vertebrates [6] and wet, high-Re aquatic swimmers [5]. In contrast, lower viscosity regimes resemble the theoretical prediction for a body elasticity dominated torque balance[6], previously not seen in experiment.

Having described the effect of the environmental forces on the phase lags, we now proceed to examine differences in muscle and curvature wave shapes and speeds in detail, and to discuss the implications of this phenomena for environmentally adaptive gait control.

### B. Environmental interactions shape undulatory waves and shorten commanded wavelengths

As noted above, phase lags can only occur if there is a concurrent difference between the phase velocity of the muscle wave v_*m*_ and the curvature wave v_*curv*_. This requires that the either the frequency f = 1/T or wavenumber k = 1/λ of the two respective waves must differ, which in turn implies that the gait commanded by the neuromuscular system (the muscle activity pattern) necessarily differs from the gait executed by the body, due to passive mechanical effects.

We find that across all body segments and environments, the frequency of the curvature and the muscle waves, as determined from fits to sine functions, are matched (Fig. 4,a). Hence, differences in phase velocity across the body arise from changes in the wavenumber. To determine the wavenumber difference, we calculated the wavenumber shift implied by a given phase lag profile, and used measured phase lags as an input.

**FIG. 4.**
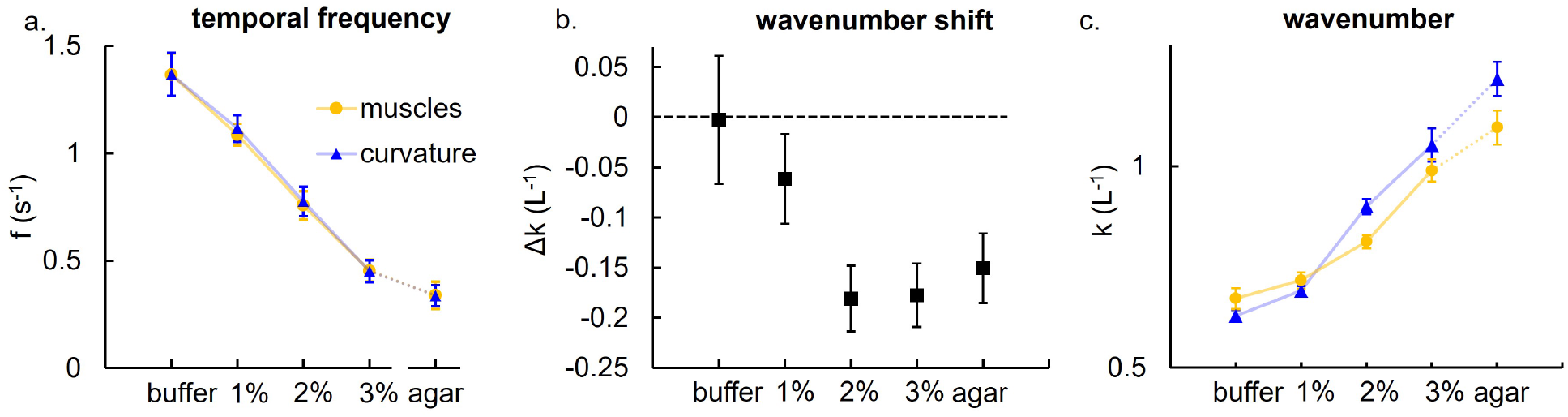
Undulatory wave parameters across the different environments. Muscle and curvature waves are frequency locked for all environments (a). However, wavenumbers shift (b) in stiffer environments (2-3% MC and agar). Absolute muscle and curvature wavelengths calculated via covariance matrices (c) show that commanded wavelengths diverge from the actual performed gait as resistance goes up, implying mechanical modulation.

The phase difference Δα between two sinusoidal traveling waves with wavenumbers k_*m*_ and k_*curv*_ and a common frequency *f* is given by

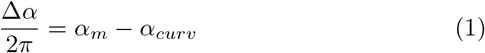

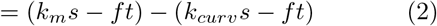

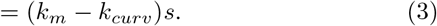

We can therefore relate the change in measured phase lags *ϕ*/*T* across a distance Δs along the body to the resultant wavenumber difference via

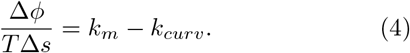

Because the phase lags do not simply increase linearly along the body, we estimated the approximate effective wavenumber shift using the average difference in phase across a region in the middle of the body, avoiding the most anterior and posterior body segments (this helped to mitigate deviations from linearity near the head and tail). The resulting wavenumber shifts are shown in Fig. 4, b. In buffer, the wavenumber shift calculated from the average slope is zero. As the drag forces increase and the phase lags begin to accumulate in the posterior region, the predicted wavenumber shift increases.

We also calculated the wavenumbers directly from the curvature and muscle activity patterns using a covariance based approach. To calculate the frequencies, the fluctuations of the muscle and curvature in time observed at a specific point could be fit reliably to sinusoids or analyzed spectrally. In contrast, for spatial variation in muscle and curvature, the finite length of the worm, the presence of noise in the muscle pattern and the deviations from sinusoidal idealization caused these approaches to become unreliable. The absolute wavenumbers were therefore calculated from covariance-matrices (see methods). These wavenumbers show a divergence between muscle and curvature at higher environmental resistances (2-3% MC and agar) in accord with the calculation derived from measured phase lags (Fig. 4, b), as the overall value of both muscle and curvature wavenumbers increase with viscosity. For lower resistances, such as buffer, rather than observing wavenumbers that converge, corresponding with a wavenumber shift of zero (as observed above in Fig. 4, b), we instead observed a wavenumber shift in the opposite direction, such that the muscle wavenumber is larger than the corresponding curvature wavenumber. This discrepancy likely reflects the region of negative slope in the phase lags in the anterior half of the organism, which corresponds with a local effective wavenumber shift in the positive direction (a phase advance).

Beyond wavenumber shifts, the non-uniformity of the phase lags along the body imply differences in the overall shape of muscle and curvature waves and different overall deviations from the sinusoidal idealization. While many approaches approximate both the body kinematics and muscle activity patterns as traveling sine waves[6], the measured waves display significant deviations from sinusoidal shapes.

To describe the average shape across time and populations, we computed population-averaged covariance matrices for both muscle and curvature waves, shown in Fig. 5, left and center columns. The width of the central lobe along the diagonal is proportional to the wave speed (and therefore the effective wavenumbers shown in Fig. 4, c), and can hence be used to visualize the variation of the speed along the body.

**FIG. 5.**
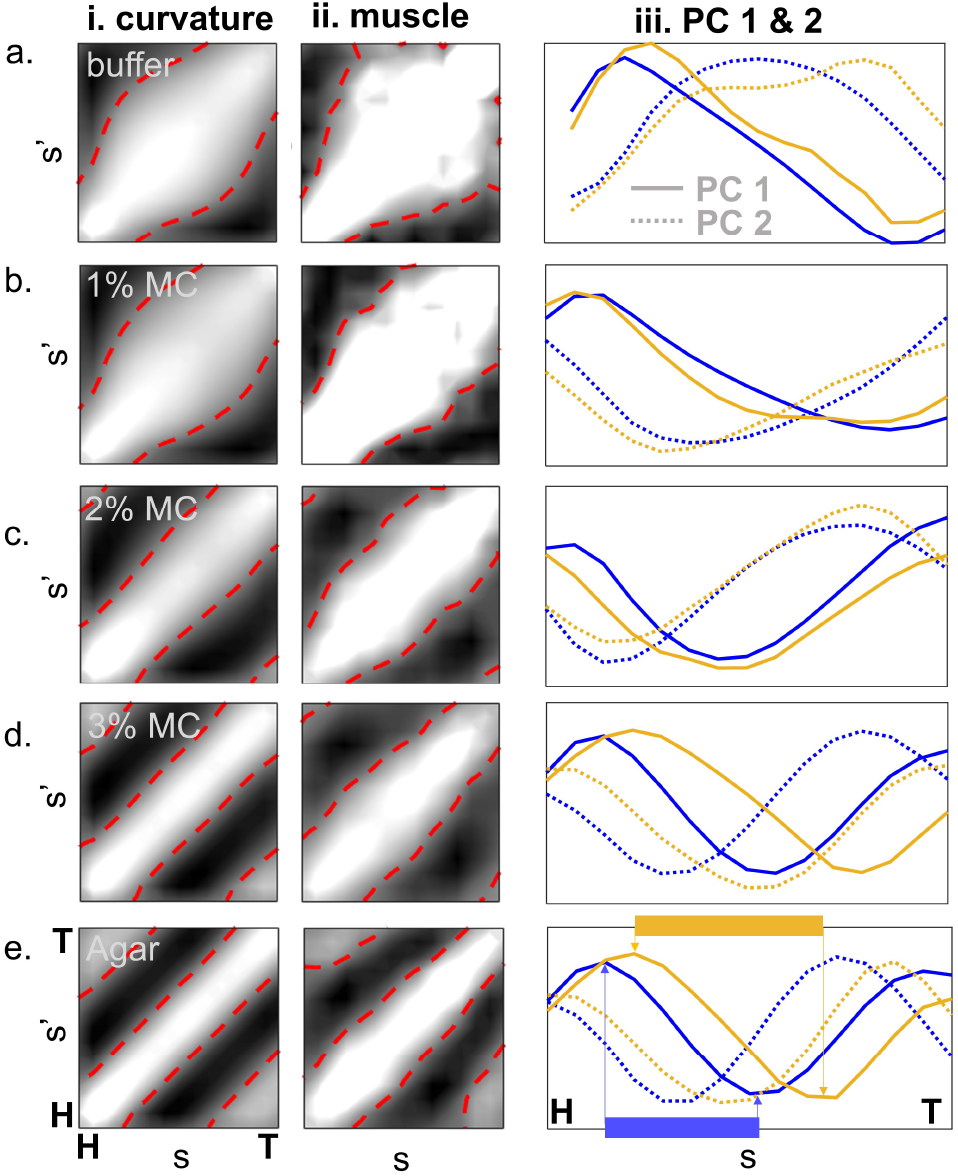
Wave shape differences account for observed phase lags. Population and time-averaged covariance matrices for curvature waves (i) and muscle activity waves (ii) and corresponding PCs 1 and 2 (iii.) for curvature (blue) and muscle activity (yellow). Panels a-e show the same quantities for buffer, 1% MC, 2% MC, 3% MC and agar, respectively.

In lower resistance environments, the variation of wave speed along the body in both the muscle and curvature waves increases until the midpoint of the body, before tapering back to a slower speed in the posterior region of the worm. As the resistance of the medium is increased, the speed variation in the curvature wave is reduced relative to the corresponding muscle wave. (See for example the relative constancy of the node-lines in the agar covariance matrices relative to the corresponding matrix for the muscles, Fig. 5, e).

We can further visualize differences in wave shape (and wavenumber), by examining the first two principle components of each respective wave (the first two eigenvectors, or “eigenworms” [26] of the covariance matrices). Fig. 5, e, highlights the peak-trough distance in the muscle waves, illustrating the wavenumber difference discussed above reflected in the shapes of the PCs. Taken together, these results indicate external forces not only induce wavelength shifts, they also shape the wave commanded by the nervous system, rectifying wavespeed variations along the body to produce a wave that more closely resembles a traveling sine wave.

### C. The relative strength of viscosity and body elasticity controls passive mechanical wave modulation

We have shown that phase lags accumulate along the body in higher resistance environments, and self-consistently, that wavenumber and ultimately wave speed differences produce the observed phase lags. Previous models predicted that body elastic torques are *in phase with body curvature* [6] and hence produced negligible phase lag accumulation. In contrast, large external torques (e.g. viscous) were predicted to produce phase lag accumulation. We, therefore, hypothesized that our experimental observations arise because internal elastic torques dominate in these environments, and that our observed phase lag accumulation in higher viscosities and agar arise from external drag dominance. To test this, we modified a previous model based on torque balance [6], that also accounts for how changes in wavelength[27] effect the phase lags, to systematically test the effect of different elastic/viscous torque ratios on the predicted phase lags.

We first constructed a model that takes observed kinematics as an input, and calculates the resultant torques from fluid drag and body elasticity, using measured passive body elasticity values for *C. elegans [*12] and measured values of the elasticity of each methylcellulose environment as parameters (See Methods). We then used torque balance to infer the muscle torque needed to realize the input kinematics as in [6]. Phase lags were then calculated by comparing the input kinematics to the inferred muscle pattern. Figure 6, a, shows that this “backward” kinematic model reproduces the experimentally measured phase lag profiles and captures the effect of increasing viscosity. This suggests that the relative magnitude of internal and external forces controls the onset of phase lag accumulation with viscosity. While previously measured phase lags were observed solely in organisms occupying the external-drag-dominated regime, *C. elegans* interpolates between the two regimes as it moves through increasingly viscous media.

**FIG. 6.**
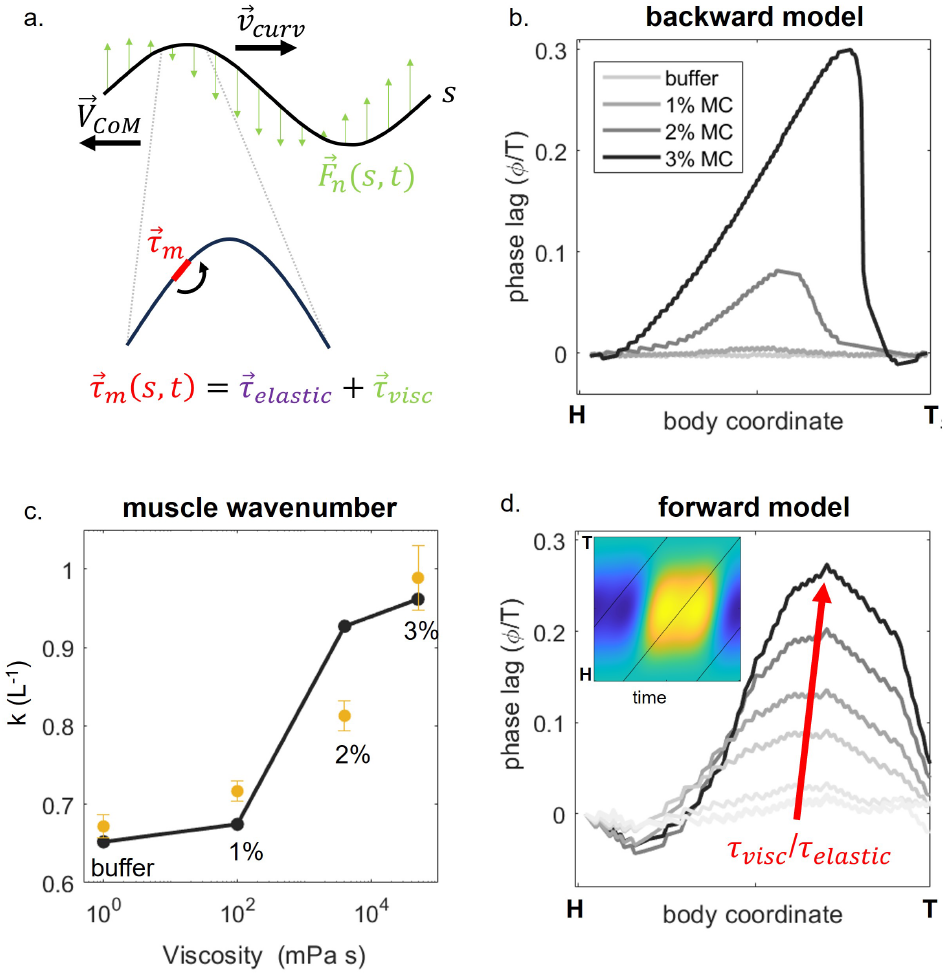
Comparison of model and experimental results. a) We model the worm as a thin elastic rod performing a sinusoidal gait. In the “backwards model”, we calculate the forces and torques due to elasticity and due to viscous forces, using a resistive force theory to determine the muscle torque pattern. b) The “backwards” model reproduces experimental phase lags using kinematics as an input. c) The “backward model also predicts muscle wavenumber trends d) In our open loop”forward” model, a muscle pattern is taken as an input and the kinematics are calculated. This model reproduces phase lag behavior with no neuronal gait modulation.

In addition to the phase lags themselves, we also used our “backward model” to calculate the wavenumber of the muscle wave and find reasonable agreement to experimental values (See Fig. 6,c). This shows that in our model, as well as in experiments, the phase lags co-occur with wavelength shifts, implying that in cases where phase lags are large, there exists a significant discrepancy between the commanded wavenumber and the wavenumber of the gait that results.

Finally, we asked whether phase lags could, in principle, arise solely from passive effects. To do so, we ran our mechanical model ‘forwards.’ We started with a single muscle pattern, obtained from the ‘backward’ model’s prediction of the muscle pattern for buffer. Using this muscle pattern as our sole input, we then varied the strength of viscous and elastic torques, and solved the ‘forward’ problem to produce the kinematics that result. We found that similar phase lag profiles may be obtained (see 6, c), implying that simply tuning the viscosity, in the absence of active changes to the wavenumber of the muscle pattern can in principle produce phase lags and the corresponding wavenumber modulation entirely through passive mechanics. While *C. elegans* does not appear to make use of a purely passive strategy, these results suggest that viscous dependent phase lags and wavenumber modulation are a generic feature of undulation that do not require active wavenumber shifts to accompany changes in viscosity.

## III. DISCUSSION AND CONCLUSION

Fundamentally, phase lags arise because, for any slender undulator, the muscle acting at a point s must balance not only torques originating from nearby body segments but also from distant points along body – simply because the torque acting at a point s is a sum of torques from point forces acting at all points between s and the end of the body [6]. In this way, the neuromechanics of undulation differs from certain limbed systems with small numbers of joints – the muscle acting on a particular joint may be insensitive to the activity of far away joints and other limbs. For undulators, the muscle acting at a point s balances a sum of torques that arise both from fundamentally different forces (e.g. external fluid drag, the internal forces from body viscoelasticity), but also acting at points along the entire body.

These different torque contributions can be either in phase (their maxima coincide with the max curvature) or out of phase of the bending wave (their maxima are mis-aligned with peak curvature). Hence, their relative strength determines whether and to what degree muscle activity coincides with the curvature profile along the body. Specifically, a previous analytical calculation has shown that external drag (e.g. granular or viscous) produces torques that are typically out of phase with curvature while the internal torques arising from body elasticity, for example, is in phase[6]. Hence, the relative strength of internal and external passive torques, as set by the mechanical properties of tissues and the surrounding environment determine the size and spatial variation in the phase lags.

We have illustrated this effect experimentally using nematodes, finding that as environmental drag is increased relative to body elasticity, neuromechanical phase lags begin to accumulate along the body. This observation is consistent with the predictions of neuromechanical models of nematodes crawling on agar[13, 18, 19] and also with a simple, mechanical developed to describe vertebrate undulation[6, 27].

In contrast to measurements of NPL via EMG recordings, the spatial resolution afforded by an optical approach has enabled us to compare the waveshapes of muscle and curvature waves. Pierce-Shimamora et al[28] compared muscle activity patterns in buffer and agar and found substantial differences; however, their reported muscle patterns did not appear to take the form of a smoothly traveling wave of activity, making determinations of phase lags and the wavelengths of muscle activity patterns prohibitively difficult. Butler et al[20] measured muscle activity patterns in a broad range of viscosities and on agar, and found that the average phase lag across the entire body remained consistant across environments, but did not report the spatial profile of the phase lags across environments. Both of these studies employed young adult worms, which we found produced inconsistent optical signals, possibly due to the variable extent of agar depression resulting from movement, the size (diameter) of the animals, the presence of vulva muscles and inconsistencies of mysoin expression after development. In this work, by instead employing larval worms at the L3 stage, we found consistent wavelike activity.

These results showed that phase lags arise due to differences in wave speed variation along the body and corresponding shifts in the wavenumber between neuronally commanded muscle activity patterns, and the curvature waves that result. We, therefore, concluded that environment-dependent wavelength tuning in *C. elegans* is not solely controlled by neural feedback, but through a combination of neural and passive mechanical effects. The passive mechanical wavenumber shifts are controlled by the elasticity of the body relative to the environmental drag.

For other undulators, including previously studied vertebrate systems, this means the NPL also produces a wavenumber discrepancy between muscle and body curvature waves, and that in general, torque balance in the majority of environments requires modulation of the wave pattern commanded by the nervous system. This is important in highly dissipative, non-inertial environments, such as low-Re fluids, agar gels, or the movement of a sandfish in sand (a frictional fluid), because the lack of inertia means that performance metrics depend solely on geometry. In low-inertia situations, the wavelength and amplitude fully determine the performance of a particular gait (for example the wave efficiency). Thus, wavenumber selection is an important determinant of performance.

Moreover, resistive force theory calculations show that wave efficiency is a non-trivial function of the wavenumber, leading to optimally performing wavelengths that change with environmental parameters, such as drag anisotropy. This explains, in part, why *C. elegans* selects higher wavenumbers as environmental resistance increases. While viscous fluids typically produce a drag anisotropy of ∼ 1.4[29], higher resistance environments, like agar surfaces, have produced higher measured drag anisotropies, estimated to be ≈ 110 [13]. Fig. 7 shows the relationship between wave efficiency and wavenumber for a viscous drag anisotropy and for estimated agar drag anisotropy. As the drag anisotropy increases, the most mechanically efficient spatial frequencies (1/λ) are larger, corresponding with greater efficiency at shorter wavelengths. Because the effect of mechanics in more resistive environments is to shift wavenumbers upward relative to the muscle wavenumber, these passive mechanical effects are adaptive, meaning that achieving a better wavelength with passive effects would reduce the requirement of larger control changes from the nervous system.

**FIG. 7.**
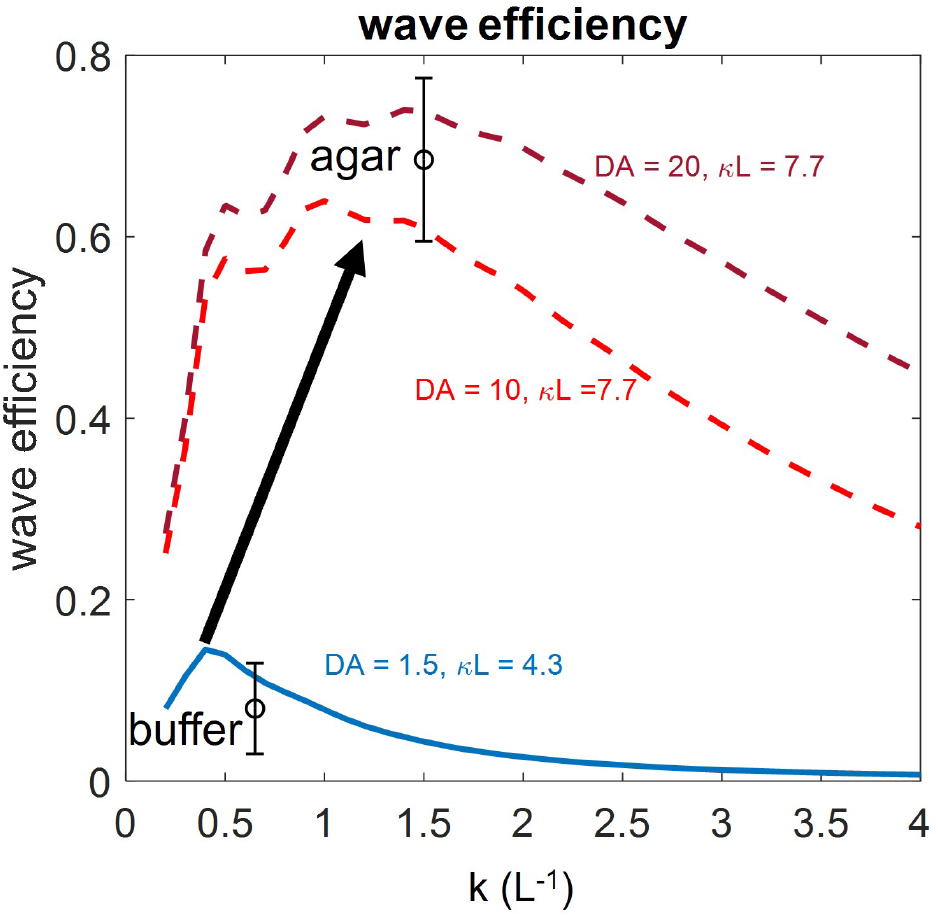
Higher drag anisotropies select for higher spatial frequencies. RFT prediction for wave efficiency for agar surfaces (dashed lines) and buffer (blue lines) along with observed spatial frequencies

Beyond comparisons of buffer and agar, similar arguments may apply to the methylcellulose experiments. While an ideal Newtonian fluid should have a drag anisotropy of 2 for all viscosities, MC solutions increase viscosity through the addition of long, polymer molecules, which are likely to increase drag anisotropy relative to the ideal Newtonian case. In general, if drag forces are accompanied by an increase in the drag anisotropy, as is likely the case in many non-newtonian biological fluids, the passive mechanical wave modulation is adaptive.

Figure 8 summarizes our model of the role of mechanics in nematode gait adaptation. As the strength of viscous environmental drag is increased, passive mechanics as well as mechanosensory neural feedback work together to select an adaptive gait. Future descriptions of nematode locomotion control should consider effects outside the nervous system. Beyond nematodes, NPL likely causes changes between commanded and actual gait wave parameters. While most previously measured examples of NPL appear to be in the high drag regime of nematodes (with negligible body elasticity relative to the muscle torque), other organisms where body elastic torques and environmental torques are comparable may display similar shifts in NPL spatial patterns upon environmental changes. These results may also suggest a new paradigm for gait selection in undulatory robots, if the mechanics of robot-environment interactions [17] could be appropriately tuned to leverage the mechanical control scheme employed by nematodes, generalized to other, complex terradynamic terrains.

**FIG. 8.**
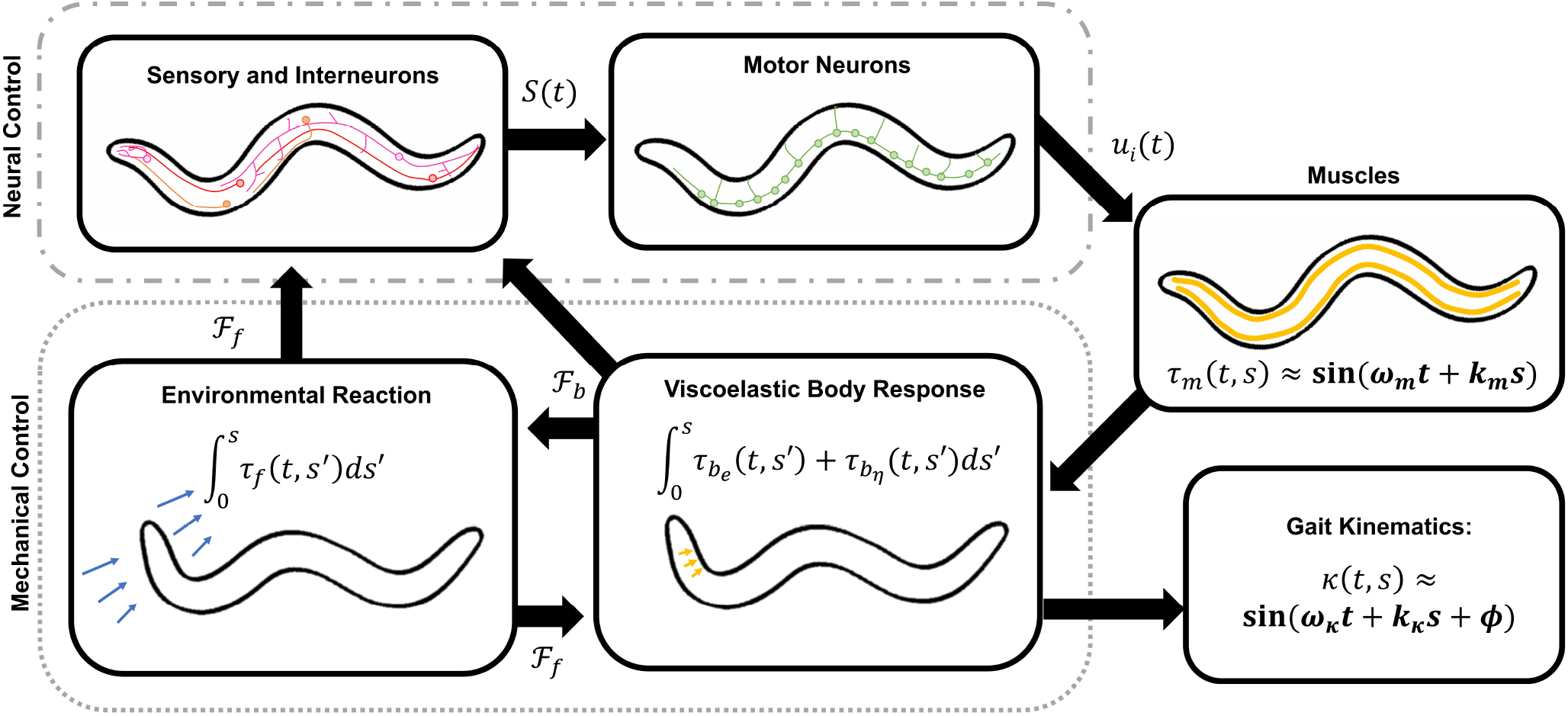
Neuromechanical system diagram for nematode gait adaptation. Mehanosensory neurons response to external physical forces and pass signals to interneurons which integrate and process the signal before sending commands to the motor neurons. Local mechanosensory signaling in the form of curvature sensing proprioception *P* (*t*) is thought to modulate motor neuron activity directly. Motor neurons pass commands to cotnract/relax downstream to muscles, which respond by creating a torque wave *τ*_*m*_(*s, t*). The torque wave is then modulated by the viscoelastic body forces and external environmental reaction forces to produce the final gait kinematics. Both mechanical and neural control interact to perform environmentally dependent gait adaptation.

## IV. METHODS

### A. Strains

Strains were maintained at 20°C and were fed *E. coli* (OP-50) bacteria on agar plates using standard protocols[30]. The transgenic lines AQ2953 ljIs131[Pmyo-3::GCaMP3-SL2-tagRFP-T] and AQ2954 ljIs132 [Pmyo-3::GFP-SL2-tagRFP-T] [20] were used to perform calcium imaging

### B. Calcium Imaging

Calcium imaging experiments were performed on a standard wide-field epifluorescence microscope described previously in[31] Synchronized worms in stage L3 (see Results above for discussion of the use of stage L3) were placed either on an agarose pad or into a small droplet of fluid between two coverslips separated by a kapton tape spacer. Fluids used were S-basal buffer solution with varying weight percentages of methylcellulose (from 0-3%) to provide a range of viscosities.

### C. Image Processing

Videos were manually cropped to sections containing bouts of forward locomotion without transients or reorientation behaviors and analyzed using protocols similar to [21, 32]. RFP and GCaMP channels were aligned using MATLAB’s imregister function. After registration, the red channel was used to segment the worm body using the canny edge detection algorithm to create a mask over the worm body in the coordinates of the registered images. This mask was then skeletonized to identify the centerline of the worm body. This centerline was subsequently smoothed using a B-spline function and used to calculate the curvature across the body as a function of time. The centerline was also used to define different muscle regions along the body in conjunction with the mask. A line normal to each point of the splined centerline was calculated and used to divide the worm into 100 discrete segments per side. Segments were made of quadrilaterals formed by intersecting the normal lines associated with two adjacent spline points with the edge of the mask. The intensity in each segment was calculated for each time point and then scaled by the number of pixels. Additionally, a small region with a fixed distance outside the worm mask was used to calculate local background intensity. The local background was subtracted from each segment in both the red and green channels, and finally, the ratio of green to red was computed.

In some cases, worms displayed consistent baseline shifts in the ratio along the body, to remove these shifts, the time average of the ratio was computed for each discrete body segment and subtracted off. Finally, to improve signal-to-noise in subsequent calculations, at the expense of spatial resolution, multiple adjacent segments were averaged creating larger blocks. For phase lag calculations segments were grouped into 9 segment blocks spanning from head to tail, and for wave shape calculations requiring higher spatial resolution segments were grouped into 15 blocks.

### C. Phase Lag Calculation

To compute phase lags, the average temporal period of the curvature wave κ(s_*i*_, t) was first computed for each segment block by first calculating the time-time autocorrelation for a given block i, and averaging the distance between adjacent peaks. Then, the phase lags were computed by calculating the cross-correlation of the muscle activity with the curvature, determining the shift of the central maximum from zero, and scaling by the temporal period of the curvature wave. To average over both body wall muscles, the left side relative to the direction of crawling was inverted before calculating the cross-correlation. m_*l*_(s_*i*_, t) to eliminate the contralteral phase shift of 180 degrees. This was performed for N = 9 segments moving from head to tail.

### E. Wavenumber Calculation

The wavenumbers were calculated using the spatial covariance matrices of curvature and muscle activity respectively (See Fig. 5, left and middle columns). By determining the position of the line of zero-crossings, we can infer the distance separating body segments oscillating ninety degrees out of phase, and therfore calculate the phase velocity. The effective wavenumber was then computed by dividing the phase velocity by the measured frequencies by the corresdponding phase velocity to produce the wavenumber.

### F. Numerical model

We used two kinds of models to compute the relative phase between the curvature and the torque generated by the muscles. In both models, the body of the nematode was considered as a uniform cylinder with a length of 1.1 mm. The overall motion of the nematode is computed using the classic resistive force theory (RFT)[33]. In RFT, the body of the nematode was divided into infinitesimal segments, and the forces F were decomposed into forces normal to the axis (midline) of the cylindrical body *F*_*n*_ and the forces parallel to the axis *F*_*l*_. The forces were proportional to the respective velocity components, i.e. *F*_*n*_ = *C*_*n*_v_*n*_ and *F*_*l*_ = *C*_*l*_*v*_*l*_. The drag coefficient in the normal direction was taken as *C*_*n*_ = 3.3η, where η is the viscosity of the fluid. The ratio between the drag coefficients *DA* = *C*_*n*_/*C*_*l*_ characterizes the anisotropy of the fluid. The motion of nematode was constrained in a plane, and the three degrees of freedom were considered. Assuming the inertia is negligible, force (torque) balance was reached at every moment. Then the translational and rotational velocities were obtained. Since the anisotropy is unknown and the swimming speed was measured, we tuned DA to match the swimming speed. Although the anisotropy has a significant effect on swimming speed, it affects the pattern of the torque little. The differences between the wavelengths of the torque generated by a constant anisotropy DA = 1.6 and speed-matched values is less than 5%.

In the “backward” model, the kinematics was prescribed as a traveling wave *κ*(*s, t*) = *A*_*κ*_ sin(2*πft* + 2*πs*/*λ*), where *κ* is the curvature of the midline of the body, *s* is the arc length along the midline, *A*_*κ*_ is the amplitude, *f* is the frequency, *t* is time, and *λ* is the wavelength. These kinematic parameters were set as those values from the experiments. After the motion was solved, we computed the torque based on the force distribution along the body as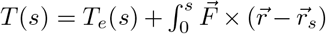. The torque contribution from the body elasticity was *T*_*e*_(*s*) = *bκ*(*s*). The elasticity of nematode body had been measured previously in several studies, but the values spanned a few orders of magnitude and the values were obtained with anesthetized worms[12, 34, 35]. Nonetheless, it is safe to say that the torque from elasticity is significant when nematodes swim in low viscosity environments. Therefore, we varied the body elasticity b in the range of 2 − 200 *×* 10^14^*Nm*^3^. The torque contribution from the body viscosity is negligible and hence not considered in our model, according to previous study [12].

In the open-loop (“forward”) model, the muscle torque pattern was fixed and prescribed as a traveling wave pattern (See Supplementary Figure SXX for the constructed torque pattern). This torque pattern was constructed from a “backward” model, in which the wavenumber of 1.5 and the body elasticity is tuned to slightly change the torque near the tail and decay near head. The torque was scaled by a factor in each case such that the resulted average amplitude of the motion matched the amplitude at the same viscosity in experiment. In the open-loop model, the elasticity of the body was fixed to 4.5 *×* 10^−12^*Nm*^3^. To stably solve the deformation (curvature) of the body, we first assumed that at every point of the midline the curvature was a sinusoidal function of time, i.e. *κ*(*s, t*) = *A*_*B*_(*s*) sin(2*πft* + *ϕ(s)*), where the *A*_*B*_(*s*) is the amplitude of the curvature, *f* is the frequency and the value is the same as the input torque, *ϕ*(*s*) is the phase. We took equally spaced 11 points of *s*_*i*_ and used *A*(*s*_*i*_) and *ϕ*(*s*_*i*_) as the unknown variables for solving the motion. *A*_*B*_ and ϕ for any value of s was then linearly interpolated by those corresponding 11 points. Due to the imposed functional form of the curvature, input torque can not be perfectly matched, and the magnitude of the residual torque was ≈5% of the magnitude of the input torque. The resulted phase ϕ(s_*i*_) sometimes was not monotonic near the ends. Therefore, we estimated the spatial frequency by a linear fit of *ϕ*(*s*) ≈ *k*_*ϕ*_*s*, where 0.05 < *s* < 0.95 and *k*_*ϕ*_ is the fitting parameter. Then the wavelength of the curvature wave was computed as 1/*k*_*ϕ*_.

## References

[1] N. Cohen and J. H. Boyle, Swimming at low reynolds number: a beginners guide to undulatory locomotion, Contemporary Physics 51, 103 (2010).

[2] S. S. Sharpe, Y. Ding, and D. I. Goldman, enEnvironmental interaction influences muscle activation strategy during sand-swimming in the sandfish lizard scincus scincus, J. Exp. Biol. 216, 260 (2013).

[3] G. V. Lauder and E. D. Tytell, Hydrodynamics of undulatory propulsion, in Fish Physiology, Vol. 23 (Academic Press, 2005) pp. 425–468.

[4] K. D’Août, N. A. Curtin, T. L. Williams, and P. Aerts, enMechanical properties of red and white swimming muscles as a function of the position along the body of the eel anguilla anguilla, J. Exp. Biol. 204, 2221 (2001).

[5] T. McMillen, T. Williams, and P. Holmes, enNonlinear muscles, passive viscoelasticity and body taper conspire to create neuromechanical phase lags in anguilliform swimmers, PLoS Comput. Biol. 4, e1000157 (2008).

[6] Y. Ding, S. S. Sharpe, K. Wiesenfeld, and D. I. Goldman, Emergence of the advancing neuromechanical phase in a resistive force dominated medium, Proceedings of the National Academy of Sciences 110, 10123 (2013).

[7] B. Jayne and G. Lauder, enRed muscle motor patterns during steady swimming in largemouth bass: effects of speed and correlations with axial kinematics, J. Exp. Biol. 198, 1575 (1995).

[8] T. McMillen, T. Williams, and P. Holmes, Nonlinear muscles, passive viscoelasticity and body taper conspire to create neuromechanical phase lags in anguilliform swimmers, PLoS computational biology 4, e1000157 (2008).

[9] B. C. Jayne and G. V. Lauder, Red muscle motor patterns during steady swimming in largemouth bass: effects of speed and correlations with axial kinematics, Journal of Experimental Biology 198, 1575 (1995).

[10] M.-A. Félix and C. Braendle, The natural history of caenorhabditis elegans, Current biology 20, R965 (2010).

[11] S. Berri, J. H. Boyle, M. Tassieri, I. A. Hope, and N. Cohen, Forward locomotion of the nematode c. elegans is achieved through modulation of a single gait, HFSP journal.

[12] C. Fang-Yen, M. Wyart, J. Xie, R. Kawai, T. Kodger, S. Chen, Q. Wen, and A. D. Samuel, Biomechanical analysis of gait adaptation in the nematode caenorhabditis elegans, Proceedings of the National Academy of Sciences 107, 20323 (2010).

[13] X. Shen and P. E. Arratia, Undulatory swimming in viscoelastic fluids, Physical review letters 106, 208101 (2011).

[14] G. Juarez, K. Lu, J. Sznitman, and P. E. Arratia, Motility of small nematodes in wet granular media, Europhysics Letters 92, 44002 (2010).

[15] S. Park, H. Hwang, S.-W. Nam, F. Martinez, R. H. Austin, and W. S. Ryu, Enhanced caenorhabditis elegans locomotion in a structured microfluidic environment, PloS one 3, e2550 (2008).

[16] T. Majmudar, E. E. Keaveny, J. Zhang, and M. J. Shelley, Experiments and theory of undulatory locomotion in a simple structured medium, Journal of the Royal Society Interface 9, 1809 (2012).

[17] T. Wang, C. Pierce, V. Kojouharov, B. Chong, K. Diaz, H. Lu, and D. I. Goldman, enMechanical intelligence simplifies control in terrestrial limbless locomotion, Sci Robot 8, eadi2243 (2023).

[18] J. E. Denham, T. Ranner, and N. Cohen, Signatures of proprioceptive control in caenorhabditis elegans locomotion, Philosophical Transactions of the Royal Society B: Biological Sciences 373, 20180208 (2018).

[19] J. E. Denham, T. Ranner, and N. Cohen, Neuromechanical phase lag predicts material and neural control properties in caenorhabditis elegans, bioRxiv, 312389 (2018).

[20] V. J. Butler, R. Branicky, E. Yemini, J. F. Liewald, A. Gottschalk, R. A. Kerr, D. B. Chklovskii, and W. R. Schafer, A consistent muscle activation strategy underlies crawling and swimming in caenorhabditis elegans, Journal of the Royal Society Interface 12, 20140963 (2015).

[21] D. Porto, Y. Matsunaga, B. Franke, R. M. Williams, H. Qadota, O. Mayans, G. M. Benian, and H. Lu, enConformational changes in twitchin kinase in vivo revealed by FRET imaging of freely moving c. elegans, Elife 10 (2021).

[22] J. Ding, L. Peng, S. Moon, H. J. Lee, D. S. Patel, and H. Lu, An expanded gcamp reporter toolkit for functional imaging in caenorhabditis elegans, G3: Genes, Genomes, Genetics 13, jkad183 (2023).

[23] M. S. Creamer, K. S. Chen, A. M. Leifer, and J. W. Pillow, enCorrecting motion induced fluorescence artifacts in two-channel neural imaging, PLoS Comput. Biol. 18, e1010421 (2022).

[24] R. A. Stiernagle, in Wormbook, edited by T. C. elegans Research Community (Wormbook, 2006) Chap. Imaging the activity of neurons and muscles.

[25] Q. Wen, M. D. Po, E. Hulme, S. Chen, X. Liu, S. W. Kwok, M. Gershow, A. M. Leifer, V. Butler, C. Fang-Yen, T. Kawano, W. R. Schafer, G. Whitesides, M. Wyart, D. B. Chklovskii, M. Zhen, and A. D. T. Samuel, enProprioceptive coupling within motor neurons drives c. elegans forward locomotion, Neuron 76, 750 (2012).

[26] G. J. Stephens, B. Johnson-Kerner, W. Bialek, and W. S. Ryu, enDimensionality and dynamics in the behavior of c. elegans, PLoS Comput. Biol. 4, e1000028 (2008).

[27] T. Ming and Y. Ding, enTransition and formation of the torque pattern of undulatory locomotion in resistive force dominated media, Bioinspir. Biomim. 13, 046001 (2018).

[28] J. T. Pierce-Shimomura, B. L. Chen, J. J. Mun, R. Ho, R. Sarkis, and S. L. McIntire, Genetic analysis of crawling and swimming locomotory patterns in c. elegans, Proceedings of the National Academy of Sciences 105, 20982 (2008).

[29] J. Sznitman, X. Shen, R. Sznitman, and P. E. Arratia, Propulsive force measurements and flow behavior of undulatory swimmers at low reynolds number, Phys. Fluids 22, 121901 (2010).

[30] T. Stiernagle, in Wormbook, edited by T. C. elegans Research Community (Wormbook, 2006) Chap. The Maintenance of C. elegans.

[31] Y. Cho, D. A. Porto, H. Hwang, L. J. Grundy, W. R. Schafer, and H. Lu, enAutomated and controlled mechanical stimulation and functional imaging in vivo in c. elegans, Lab Chip 17, 2609 (2017).

[32] J. N. Stirman, M. M. Crane, S. J. Husson, S. Wabnig, C. Schultheis, A. Gottschalk, and H. Lu, enReal-time multimodal optical control of neurons and muscles in freely behaving caenorhabditis elegans, Nat. Methods 8, 153 (2011).

[33] E. Lauga and T. R. Powers, The hydrodynamics of swimming microorganisms, Reports on progress in physics 72, 096601 (2009).

[34] M. Backholm, W. S. Ryu, and K. Dalnoki-Veress, Viscoelastic properties of the nematode caenorhabditis elegans, a self-similar, shear-thinning worm, Proceedings of the National Academy of Sciences 110, 4528 (2013).

[35] S.-J. Park, M. B. Goodman, and B. L. Pruitt, Analysis of nematode mechanics by piezoresistive displacement clamp, Proceedings of the National Academy of Sciences 104, 17376 (2007).

